# Sex-Dependent Microbiome-Metabolome Remodeling Associated with *Bacteroides acidifaciens* Modulation of Acute Respiratory Distress Syndrome

**DOI:** 10.64898/2026.10.05.756742

**Authors:** Shanieka Staley, Carly Cooper, Stacey Oxendine, Jordan Williams, Alissa Armstrong, Kiesha Wilson

## Abstract

Acute respiratory distress syndrome (ARDS) is a severe lung disorder influenced by the interplay between the intestinal microbiome, host metabolism, and immune function. We previously found that Bacteroides acidifaciens (BA) has opposing effects on staphylococcal enterotoxin B (SEB)-induced ARDS, worsening the condition in female mice while alleviating it in males. This study aimed to examine whether these outcomes are linked to sex-dependent changes in the intestinal microbiome and gut-lung metabolome. C3H/HeJ mice of both sexes were orally colonized with Bacteroides acidifaciens before SEB exposure. Stool and bronchoalveolar lavage fluid (BALF) were collected for analysis using 16S rRNA sequencing and untargeted liquid chromatography mass spectrometry metabolomics. Our findings revealed distinct sex-dependent metabolic and microbial responses. In females, disease exacerbation correlated with elevated lipid mediators (leukotriene B4, 13-oxoODE, and 13S-HODE) alongside shifts in bile acid and redox metabolism. The intestinal microbiome showed changes in Bacteroides, Akkermansia, and Lachnospiraceae. Conversely, males showed a protective phenotype characterized by increased butyrate and alterations in sphingolipid, amino acid, redox, and nucleotide pathways. Integrated analyses identified sex-associated microbial communities and metabolomic profiles, highlighting distinct patterns of lipid and redox associations in females versus sphingolipid and short-chain fatty acid patterns in males. These results suggest that host sex influences how a commensal bacterium affects inflammatory lung injury and point to pathways for future research.

**Importance:** ARDS is a fatal inflammatory response with high mortality and limited targeted therapies. Emerging evidence has pointed to the gut-lung axis as a source of inflammation that shapes pulmonary immune responses. Yet, how individual commensal bacteria influence this axis, and how biological sex alters both the microbiome and immune response remains poorly understood. *Bacteroides acidifaciens*, drives fundamentally different intestinal microbial and metabolic states in female and male mice during ARDS, and provides some evidence of how both can impact ARDS outcomes. Females develop an environment enriched in inflammatory lipid mediators, while males exhibit a metabolic profile associated with barrier support and reduced inflammation. These findings have implications for understanding sex-based differences in ARDS outcomes and for identifying microbiome-targeted therapeutic strategies.

## Introduction

Acute respiratory distress syndrome (ARDS) is a severe inflammatory lung disorder characterized by diffuse alveolar injury, disruption of the alveolar-capillary barrier, dysregulated immune activation, and hypoxemia. Despite advances in supportive care, ARDS remains associated with substantial morbidity and mortality, and effective targeted therapies remain limited [1,2]. Increasing evidence indicates that ARDS is influenced not only by pulmonary inflammation but also by systemic interactions involving the intestinal microbiome, host metabolism, and immune function [3-5]. Through the gut-lung axis, microbial communities and their metabolites can influence epithelial barrier integrity, leukocyte responses, inflammatory signaling, oxidative stress, and tissue repair [3-5]. However, how individual commensal organisms shape these responses remains incompletely understood.

Among the intestinal microbiota, *Bacteroides acidifaciens* (BA) is a metabolically active commensal capable of influencing host immunity and metabolism. BA has been associated with beneficial effects on metabolic dysfunction and intestinal inflammation through modulation of host metabolism, epithelial integrity, microbial community structure, and immune homeostasis [6-8]. However, its biological effects appear to depend on the inflammatory and physiological context, suggesting that BA may function as a context-dependent regulator of host-microbiome interactions.

Consistent with this concept, our previous study demonstrated that BA differentially modulates staphylococcal enterotoxin B (SEB)-induced ARDS according to host sex. BA colonization attenuated disease in male mice, with reduced pulmonary inflammation and preservation of epithelial barrier function, whereas BA exacerbated disease in females, which exhibited increased lung injury, neutrophilic and Th17-associated inflammation, oxidative stress, impaired intestinal barrier integrity, and enhanced IgA responses. Inhibition of estrogen synthesis with letrozole attenuated several features of the female BA-associated phenotype, further implicating the host hormonal environment in this divergent response [9].

Although these studies established opposing immune phenotypes, the microbial and metabolic changes accompanying these responses remain undefined. BA may influence host physiology directly while also restructuring the intestinal microbial community and altering the production or transformation of bioactive metabolites. Metabolic pathways involving bile acids [10], sphingolipids [11], short-chain fatty acids [12,13], tryptophan metabolites [14], and redox regulation [15] can influence epithelial and immune function and provide potential links between microbial activity and distal pulmonary inflammation. Because both microbial community structure and host metabolism can differ according to sex, integrated microbiome and metabolome analyses may provide insight into the biological environments associated with the divergent effects of BA during ARDS.

To address these questions, the objective of the present study was to define sex-dependent microbial and metabolic changes accompanying BA modulation of SEB-induced ARDS. Using 16S rRNA sequencing and untargeted metabolomic profiling of stool and bronchoalveolar lavage fluid, we evaluated BA-associated changes within each sex, directly compared female and male responses, and integrated microbial and metabolic datasets to identify sex-associated taxon-metabolite relationships. These analyses establish a systems-level framework linking the previously observed sex-dependent effects of BA with distinct microbiome and metabolome configurations during inflammatory lung injury.

## Methods

### Experimental Design, *Bacteroides acidifaciens* Colonization, and SEB-Induced ARDS

Male and female C3H/HeJ mice (8-10 weeks old) were obtained from The Jackson Laboratory and maintained under specific pathogen-free conditions at the University of South Carolina School of Medicine animal facility on a 12-hour light/dark cycle with ad libitum access to food and water. All procedures were approved by the University of South Carolina Institutional Animal Care and Use Committee.

Mice were assigned to four experimental groups: untreated controls receiving PBS vehicle (VEH), *Bacteroides acidifaciens* (BA)-colonized mice (BA), SEB-treated mice (SEB), and BA-colonized mice subsequently challenged with SEB (BA+SEB). BA was cultured anaerobically in chopped meat broth and administered by oral gavage at 1 × 10^9 CFU in PBS; control animals received PBS. ARDS was induced using a dual-dose staphylococcal enterotoxin B (SEB) exposure model consisting of intranasal administration followed by intraperitoneal injection, as previously described [9]. Mice were euthanized 72 hours after SEB administration. Primary analyses in the main figures focused on SEB versus BA+SEB comparisons within each sex.

At euthanasia, stool was collected from cecal and colonic contents and snap-frozen in liquid nitrogen for microbiome and metabolomic analyses. Bronchoalveolar lavage fluid (BALF) was collected by tracheal cannulation and lavage with 1 mL sterile PBS, centrifuged at 10,000 × *g* for 10 min at 4°C, and supernatants were retained for metabolomic profiling.

### 16S rRNA Gene Sequencing and Microbiome Analysis

Microbial community profiling was performed using 16S rRNA gene sequencing of stool-derived bacterial DNA. Because the study was designed to evaluate microbiome changes associated with Bacteroides acidifaciens (BA) colonization, mice were maintained in treatment-specific cages rather than co-housed across experimental groups to minimize microbial transfer and preserve differences in colonization status. Housing conditions were otherwise maintained consistently across experimental groups.

DNA was extracted from stool samples using the DNeasy PowerSoil Pro Kit (Qiagen, Germantown, MD, USA) according to the manufacturer’s instructions. The bacterial 16S rRNA gene was amplified using the universal primers 515F and 806R, and amplicons were sequenced on an Illumina MiSeq platform using 2 × 250-bp paired-end sequencing.

Sequence data were processed using QIIME 2 (version 2023.9). Raw sequencing reads were quality filtered and denoised to generate amplicon sequence variants (ASVs), and taxonomic classification was performed using the SILVA 138 reference database. Downstream microbial community analyses and visualization were performed using MicrobiomeAnalyst and R-Studio. Feature-table and taxonomy data were analyzed in R. Stool samples were grouped by sex and treatment. For relative-abundance plots, features were collapsed to the genus level, converted to relative abundance, and averaged within each sex-by-treatment group. The most abundant genera were displayed in stacked bar plots.

Beta diversity was calculated using Bray-Curtis dissimilarity and visualized by principal coordinates analysis (PCoA). Shaded outlined ovals were used as visual group guides only. Sex-biased genera were explored using log2 fold changes in mean relative abundance between females and males within each treatment.

### Untargeted Metabolomic Profiling

Untargeted metabolomic profiling was performed on stool and BALF samples by General Metabolics (GMet) using liquid chromatography-mass spectrometry (LC-MS)-based metabolomics. Stool and BALF samples were extracted in 80% methanol, centrifuged to remove precipitated proteins and debris, and supernatants were analyzed in both positive and negative ionization modes. Metabolite features were identified based on accurate mass and retention time matching against internal reference databases.

Raw metabolite intensity data were normalized prior to downstream analyses and processed using MetaboAnalyst 5.0. Principal component analysis (PCA), partial least squares discriminant analysis (PLS-DA), heatmap clustering, and volcano plot analyses were performed to identify differential metabolic signatures between treatment groups. Pathway enrichment and impact analyses were conducted using Kyoto Encyclopedia of Genes and Genomes (KEGG)-associated pathways. Significant metabolic pathways were defined using FDR-adjusted p < 0.05. Integrated pathway analyses were used to identify sex-dependent metabolic remodeling patterns associated with inflammatory or protective ARDS phenotypes.

### Integrated Gut Microbiome-Metabolome Correlation Analysis

Integrated microbiome-metabolome analyses were performed using MicrobiomeAnalyst to identify functional associations between microbial taxa and metabolite abundance patterns across experimental groups. Data matrices were filtered to include significantly altered microbial taxa and metabolites (FDR < 0.05). Pairwise associations between microbial and metabolic features were calculated using Spearman’s correlation coefficient (ρ). Significant correlations were visualized as microbiome-metabolome interaction heatmaps and pathway-associated correlation networks.

Integrated enrichment analyses were used to identify interconnected microbial and metabolic pathways associated with sex-dependent responses to BA colonization during SEB-induced ARDS. These analyses were performed to define systems-level gut-lung interaction networks associated with inflammatory metabolic remodeling in females and adaptive metabolic preservation in males.

### Statistical Analysis

Statistical analyses were performed using GraphPad Prism, MetaboAnalyst, and R-based bioinformatic pipelines. Data are presented as mean ± SEM unless otherwise indicated. Group comparisons were performed using appropriate parametric or nonparametric statistical tests depending on data distribution. Statistical significance was defined as p < 0.05. Principal component analyses, heatmaps, volcano plots, diversity analyses, and pathway enrichment analyses were generated using standard metabolomic and microbiome analytical workflows.

## Results

### Experimental Framework for Sex-Dependent Microbiome-Metabolome Remodeling During BA-Associated ARDS

Our previous study demonstrated that *Bacteroides acidifaciens* (BA) exacerbates SEB-induced ARDS in female mice while attenuating disease severity in males. Female BA+SEB mice exhibited increased pulmonary inflammation, oxidative stress, neutrophil recruitment, IgA responses, and IL-17-associated signaling, whereas male BA+SEB mice exhibited reduced inflammation, improved epithelial barrier integrity, and increased IL-22-associated responses. Based on these divergent phenotypes, the present study examined whether BA-associated differences in disease severity are accompanied by sex-dependent remodeling of the intestinal microbiome and stool and pulmonary metabolomes (**Figure 1**).

**Figure 1.**
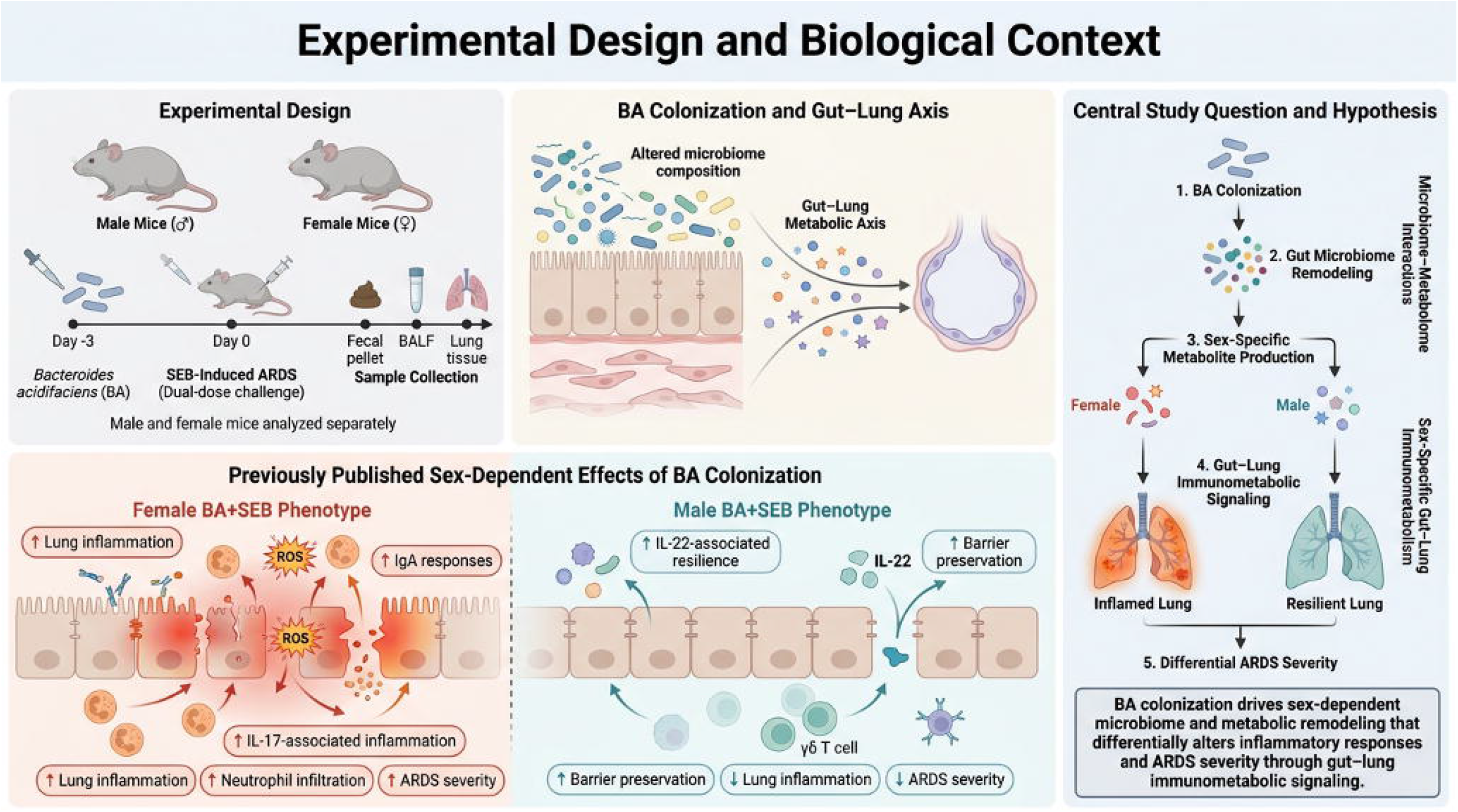
Experimental design and conceptual framework of sex-dependent microbiome-metabolome remodeling during *Bacteroides acidifaciens* (BA)-associated ARDS. Male and female C3H/HeJ mice were orally colonized with Bacteroides acidifaciens (BA) before induction of staphylococcal enterotoxin B (SEB)-induced acute respiratory distress syndrome (ARDS). Stool and bronchoalveolar lavage fluid (BALF) were collected for microbiome and metabolomic analyses. Previous studies demonstrated that BA exacerbates SEB-induced ARDS in female mice while attenuating disease severity in males. The current study examined whether these divergent inflammatory phenotypes are accompanied by sex-dependent intestinal microbiome and gut-lung metabolome remodeling.

### BA Colonization Alters Intestinal and Pulmonary Metabolomic Profiles in Female Mice During ARDS

Untargeted metabolomic profiling revealed distinct stool and BALF metabolic profiles between female SEB and BA+SEB mice. Stool PCA showed separation between treatment groups (**Figure 2A**). Cholate, aldosterone glucuronide, and indolelactic acid were increased in BA+SEB mice, whereas dopamine and betaine were decreased (**Figure 2B**). Pathway analysis identified alterations involving amino acid, sphingolipid, steroid, bile acid, linoleic acid, xenobiotic, and glutathione metabolism (**Figure 2C**).

**Figure 2.**
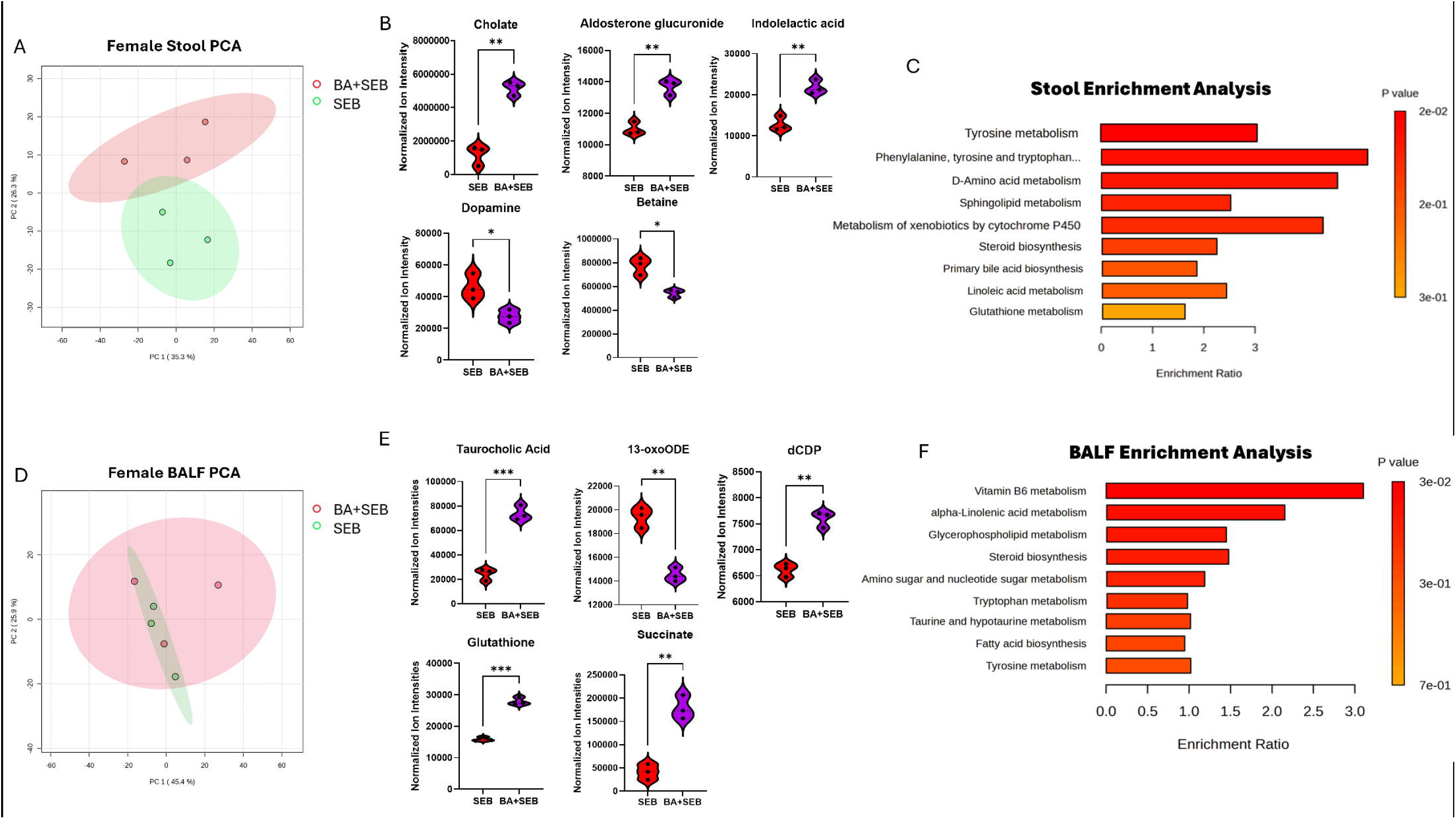
BA colonization is associated with gut and pulmonary metabolomic remodeling in female mice during SEB-induced ARDS. (A) Principal component analysis (PCA) of female stool metabolomic profiles comparing SEB and BA+SEB groups. PC1 and PC2 account for the indicated percentages of total variance. (B) Differentially abundant stool metabolites between SEB and BA+SEB mice. Selected metabolite features are shown, including cholate, aldosterone glucuronide, indolelactic acid, dopamine, and betaine. (C) Pathway enrichment analysis of female stool metabolomic profiles. (D) PCA of female BALF metabolomic profiles comparing SEB and BA+SEB groups. (E) Differentially abundant BALF metabolites, including taurocholic acid, dCDP, glutathione, succinate, and 13-oxoODE. (F) Pathway enrichment analysis of female BALF metabolomic profiles. Statistical significance is indicated as ^*^p < 0.05, ^**^p < 0.01, and ^***^p < 0.001.

BALF metabolomes also differed between female SEB and BA+SEB mice (**Figure 2D**). Taurocholic acid, dCDP, glutathione, and succinate were increased in BA+SEB mice, whereas 13-oxoODE was decreased (**Figure 2E**). Enriched pathways included lipid, steroid, amino acid, tryptophan, taurine/hypotaurine, vitamin B6, and carbohydrate-associated metabolism (**Figure 2F**). Collectively, these findings demonstrate that BA colonization induces extensive metabolic remodeling within both the intestinal and pulmonary compartments of female mice during SEB-induced ARDS.

### BA Colonization Induces Distinct Gut and Pulmonary Metabolic Remodeling in Male Mice During SEB-Induced ARDS

Untargeted metabolomic profiling revealed BA-associated remodeling in both stool and BALF from male mice during SEB-induced ARDS. Stool PCA showed partial separation between BA+SEB and SEB groups (**Figure 3A**). BA+SEB mice exhibited increased butyric acid, daidzein, eugenitin, tyramine, and ethyl tiglate and decreased dihydroceramide relative to SEB controls (**Figure 3B**). Pathway enrichment identified alterations primarily involving sphingolipid and fatty acid metabolism, amino acid metabolism, retinol and galactose metabolism, and xenobiotic processing (**Figure 3C**).

**Figure 3.**
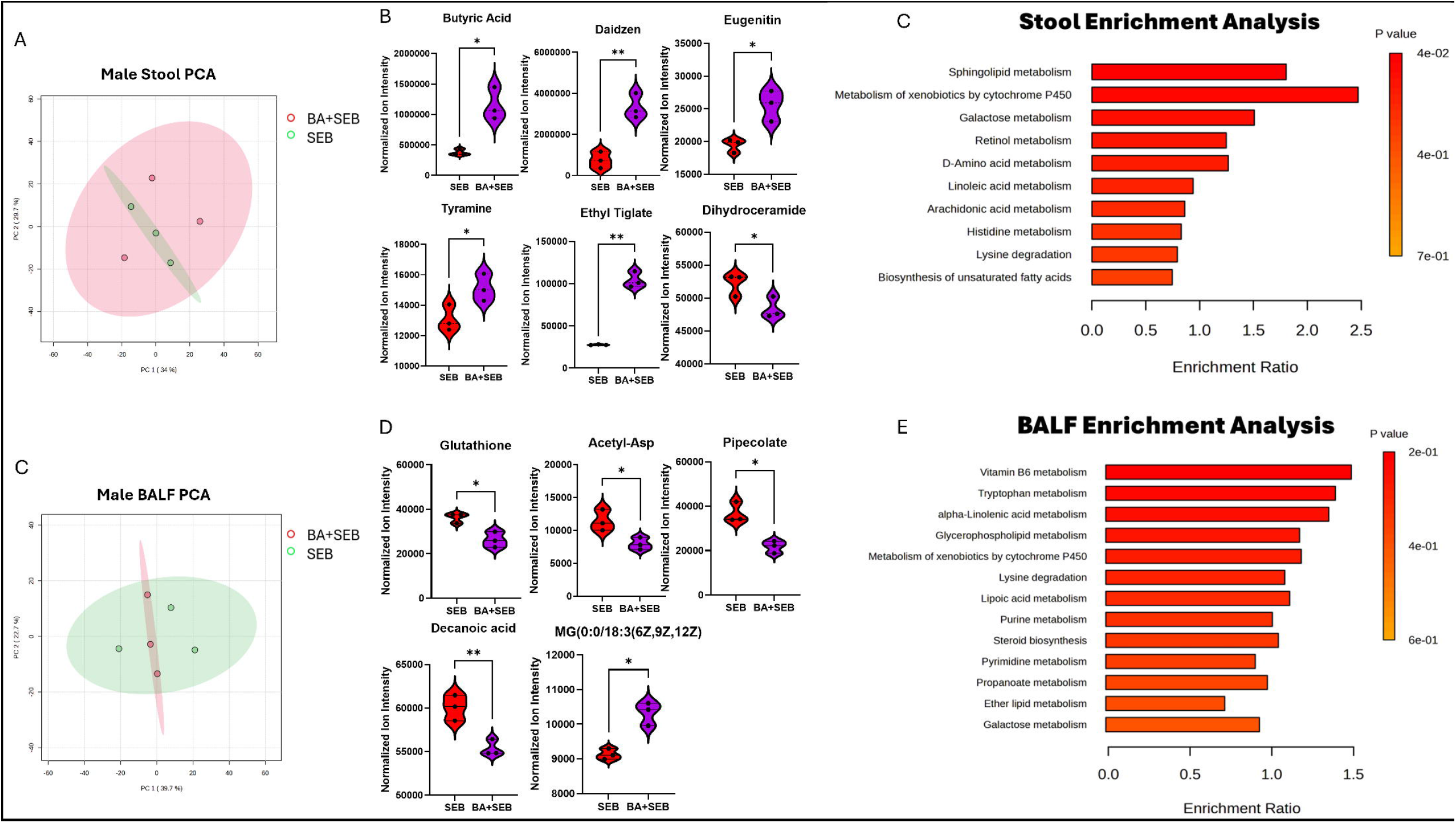
BA colonization is associated with gut and pulmonary metabolomic remodeling in male mice during SEB-induced ARDS. (A) PCA of male stool metabolomic profiles comparing SEB and BA+SEB groups. (B) Differentially abundant stool metabolites between SEB and BA+SEB mice, including butyric acid, daidzein, eugenitin, tyramine, ethyl tiglate, and dihydroceramide. (C) Pathway enrichment analysis of male stool metabolomic profiles. (D) PCA of male BALF metabolomic profiles comparing SEB and BA+SEB groups. (E) Differentially abundant BALF metabolites, including glutathione, acetyl-Asp, pipecolate, decanoic acid, and MG(0:0/18:3(6Z,9Z,12Z)). (F) Pathway enrichment analysis of male BALF metabolomic profiles. Statistical significance is indicated as shown in the individual panels.

BALF metabolomic profiles also differed between BA+SEB and SEB mice, although group separation was less pronounced than in stool (**Figure 3D**). BA+SEB mice exhibited decreased glutathione, acetyl-Asp, pipecolate, and decanoic acid and increased MG(0:0/18:3(6Z,9Z,12Z)) (**Figure 3E**). Enriched pathways included lipid, amino acid, nucleotide, vitamin B6, propanoate, and galactose metabolism (**Figure 3F**). Together, these findings demonstrate distinct intestinal and pulmonary metabolic remodeling associated with BA colonization in male mice during ARDS.

### Sex-Specific Immunometabolic Remodeling Distinguishes Female and Male BA+SEB Mice

To directly assess sex-associated metabolic differences during BA-modulated ARDS, stool and bronchoalveolar lavage fluid (BALF) metabolomic profiles were compared between female and male BA+SEB mice. Stool PCA demonstrated partial separation between sexes, indicating distinct metabolomic profiles following BA colonization and SEB challenge (**Figure 4A**).

**Figure 4.**
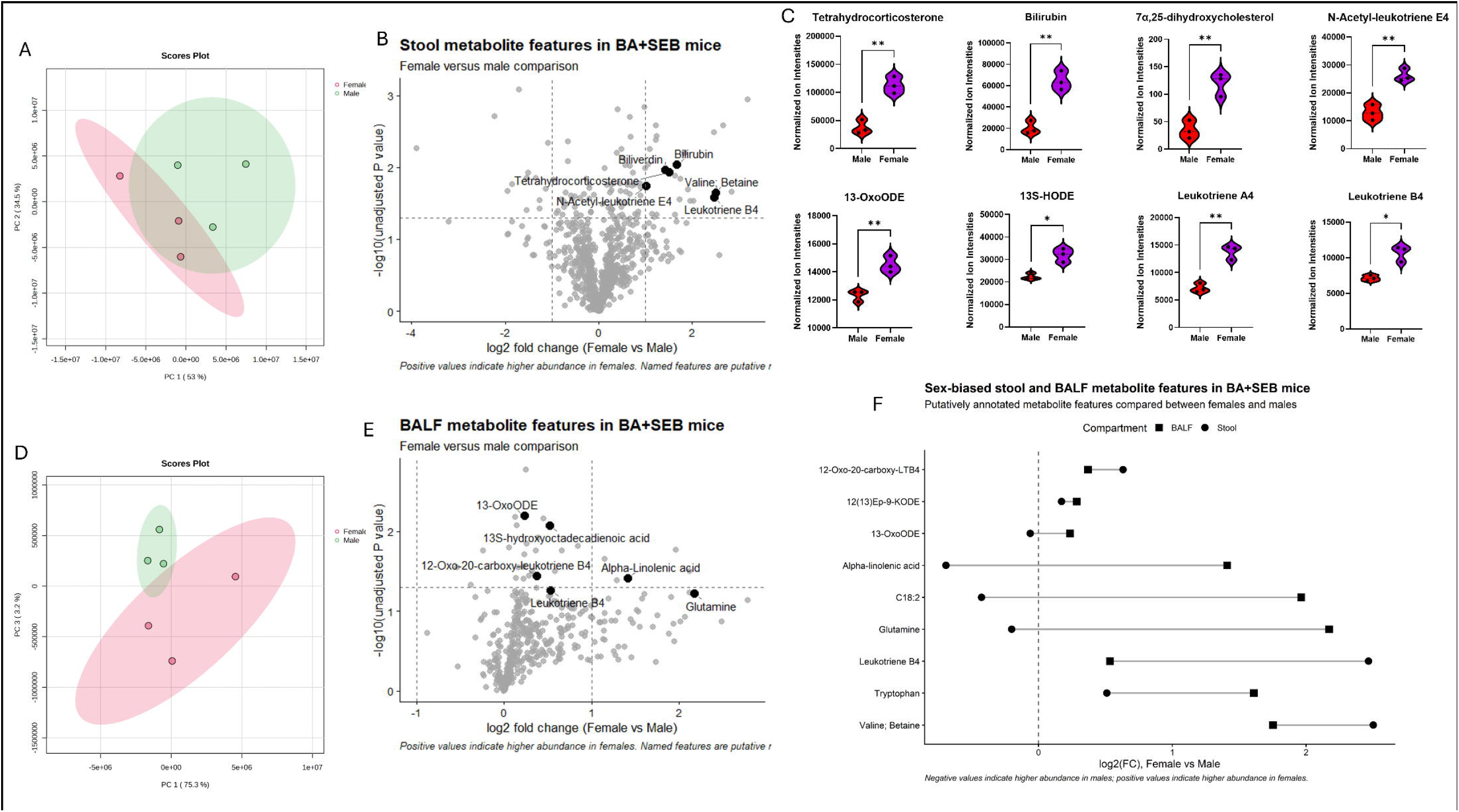
Direct comparison of stool and BALF metabolomic profiles reveals sex-associated and compartment-dependent metabolic differences in BA+SEB mice. (A) PCA of stool metabolomic profiles from female and male BA+SEB mice. Ellipses indicate group clustering, and the percentage of variance explained by each principal component is shown on the corresponding axis. (B) Volcano plot comparing stool metabolite features between female and male BA+SEB mice. Positive logL fold-change values indicate higher abundance in females, whereas negative values indicate higher abundance in males. Selected putatively annotated metabolite features are labeled. (C) Normalized ion intensities of selected stool metabolites differing between female and male BA+SEB mice. (D) PCA of BALF metabolomic profiles from female and male BA+SEB mice. (E) Volcano plot comparing BALF metabolite features between sexes, with positive values indicating higher abundance in females and negative values indicating higher abundance in males. (F) Comparison of sex-associated metabolite abundance patterns across stool and BALF. *p* < 0.05 (^*^) and *p* < 0.01 (^**^).

Differential abundance analysis identified multiple stool metabolites that differed between sexes (**Figure 4B**). Female BA+SEB mice exhibited higher levels of tetrahydrocorticosterone, bilirubin, 7α,25-dihydroxycholesterol, N-acetyl-leukotriene E4, 13-oxoODE, 13S-HODE, leukotriene A4, and leukotriene B4 relative to males (**Figure 4C**).

Sex-associated differences were also evident in the pulmonary compartment. BALF PCA showed partial separation between female and male BA+SEB mice (Figure 4D). Female mice exhibited higher abundance of several inflammatory lipid-associated metabolites, including 13-oxoODE, 13S-HODE, 12-oxo-20-carboxy-leukotriene B4, and leukotriene B4, whereas glutamine and α-linolenic acid were comparatively enriched in males (**Figure 4E**).

Comparison across stool and BALF revealed both shared and compartment-specific sex-associated metabolic patterns (**Figure 4F**). Leukotriene B4 and valine showed female-biased abundance across compartments, whereas glutamine and α-linolenic acid were relatively enriched in males. These findings demonstrate that female and male BA+SEB mice exhibit distinct metabolic states across both intestinal and pulmonary compartments.

### BA Colonization is Associated with Sex-Specific Restructuring of the Intestinal Microbiome during SEB-Induced ARDS

Microbiome profiling revealed sex-dependent changes in intestinal community structure following BA colonization during SEB-induced ARDS. In females, Chao1 richness differed across treatment groups, with BA-only mice exhibiting lower richness than VEH, SEB, and BA+SEB mice (**Figure 5A**). Bray-Curtis PCoA showed visually distinct clustering of female SEB and BA+SEB communities, indicating BA-associated restructuring of the intestinal microbiome during ARDS (**Figure 5B**). Relative abundance profiles also revealed treatment-associated shifts in dominant genera, including *Bacteroides, Akkermansia*, Lactobacillus-associated taxa, and Lachnospiraceae-associated taxa (**Figure 5C**).

**Figure 5.**
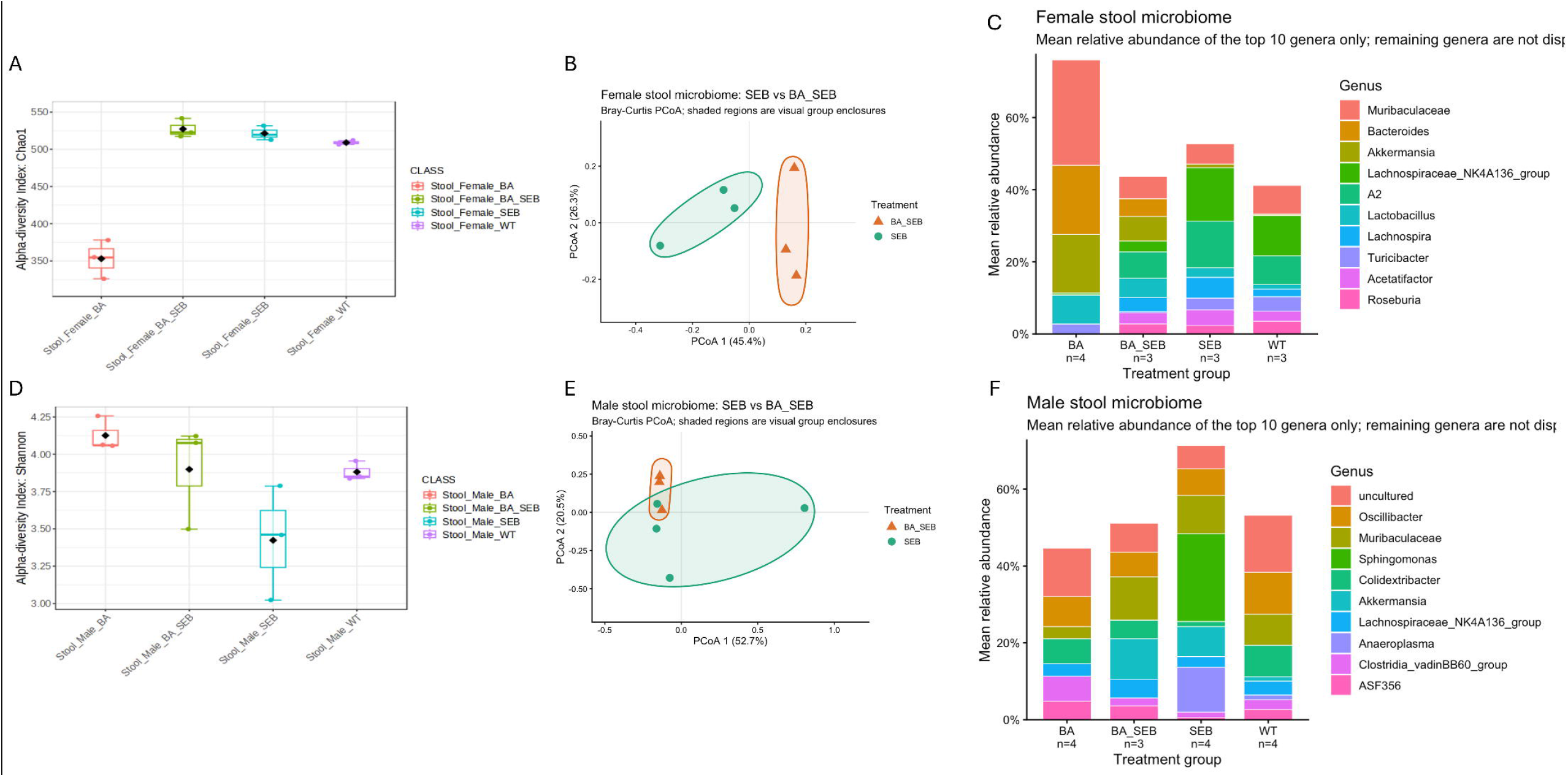
BA colonization is associated with sex-dependent restructuring of the intestinal microbiome during SEB-induced ARDS. (A) Chao1 alpha diversity of female stool microbial communities across VEH, BA, SEB, and BA+SEB treatment groups. (B) Bray-Curtis principal coordinate analysis (PCoA) of female SEB and BA+SEB stool microbiomes. Shaded ellipses indicate visual group enclosures. (C) Mean relative abundance of the dominant bacterial genera in female stool microbiomes across treatment groups. (D) Shannon alpha diversity of male stool microbial communities across VEH, BA, SEB, and BA+SEB groups. (E) Bray-Curtis PCoA of male SEB and BA+SEB stool microbiomes; shaded ellipses indicate visual group enclosures. (F) Mean relative abundance of the dominant bacterial genera in male stool microbiomes across treatment groups. Relative-abundance plots display the most abundant genera, with remaining taxa grouped or omitted as indicated in the individual panel.

In males, Shannon diversity varied across treatment groups, with SEB-treated mice exhibiting lower diversity than VEH mice and BA+SEB mice showing intermediate values (**Figure 5D**). Bray-Curtis PCoA showed greater overlap between male SEB and BA+SEB communities than observed in females, although treatment-associated differences remained apparent (**Figure 5E**). Relative abundance profiles similarly demonstrated shifts in dominant genera, including *Akkermansia*, Muribaculaceae-associated taxa, Lachnospiraceae-associated taxa, and Oscillibacter-associated taxa (**Figure 5F**).

Together, these findings indicate that BA colonization is associated with sex-dependent restructuring of the intestinal microbiome during SEB-induced ARDS. Additional analyses of selected taxa, prevalence, and BA versus BA+SEB community composition are provided in Supplementary Figures S3-S5.

### Direct Comparison Reveals Sex-Associated Differences in Intestinal Microbial Communities Following BA Colonization

Because BA produced opposing disease phenotypes in female and male mice, intestinal microbial communities were directly compared between sexes following BA colonization. Bray-Curtis PCoA showed visually distinct clustering of female and male microbiomes in BA-treated mice, and this separation remained apparent following SEB challenge (Figure 6A-B).

**Figure 6.**
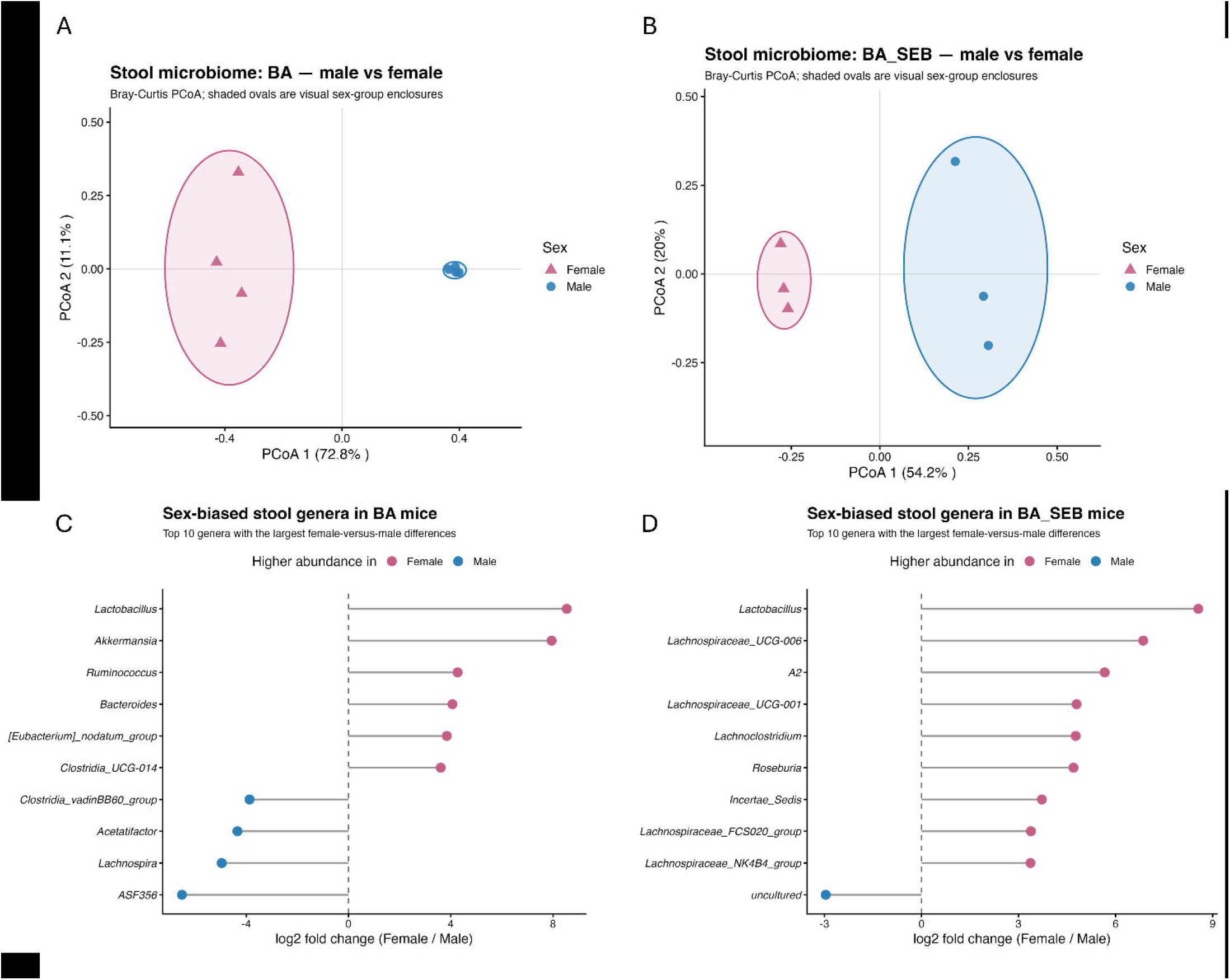
Direct comparison reveals sex-associated differences in intestinal microbial community structure following BA colonization. (A) Bray-Curtis PCoA comparing female and male stool microbiomes following BA colonization alone. (B) Bray-Curtis PCoA comparing female and male stool microbiomes following BA colonization and SEB challenge. Shaded ellipses in A-B indicate visual sex-group enclosures. (C) Genera exhibiting the largest female-versus-male differences in mean relative abundance in BA-treated mice, expressed as logL fold change (female/male). (D) Genera exhibiting the largest female-versus-male differences in mean relative abundance in BA+SEB mice. Positive logL fold-change values indicate higher mean relative abundance in females, whereas negative values indicate higher mean relative abundance in males. Panels C and D are descriptive comparisons of relative abundance and are not intended to represent formal differential-abundance testing.

Genus-level comparisons further demonstrated sex-associated differences in microbial composition. Following BA colonization alone, females showed higher relative abundance of several taxa, including *Lactobacillus, Akkermansia, Ruminococcus*, and *Bacteroides*, whereas taxa such as *Acetatifactor, Lachnospira*, ASF356, and the Clostridia vadinBB60 group were relatively more abundant in males (Figure 6C). Following SEB challenge, female BA+SEB mice showed greater relative abundance of several Lactobacillus- and Lachnospiraceae-associated taxa, including *Lachnoclostridium* and *Roseburia* (Figure 6D). These genus-level comparisons are descriptive and reflect the largest observed female-versus-male differences in mean relative abundance.

Together, these findings demonstrate that female and male mice develop distinct intestinal microbial community configurations following BA colonization and that these differences persist during SEB-induced ARDS.

### Integrated Microbiome-Metabolome Analyses Identify Sex-Associated Microbe-Metabolite Networks

Integrated microbiome-metabolome analyses identified distinct microbial-metabolic association patterns in female and male BA+SEB mice. In females, pathway network analysis highlighted coordinated changes involving amino acid, lipid, bile acid, and redox-associated metabolism (**Figure 7A**). Correlation analysis further identified broad associations between stool metabolite features and taxa including uncultured *Clostridium, Acetatifactor muris*, and *Staphylococcus carnosus* (**Figure 7B**).

**Figure 7.**
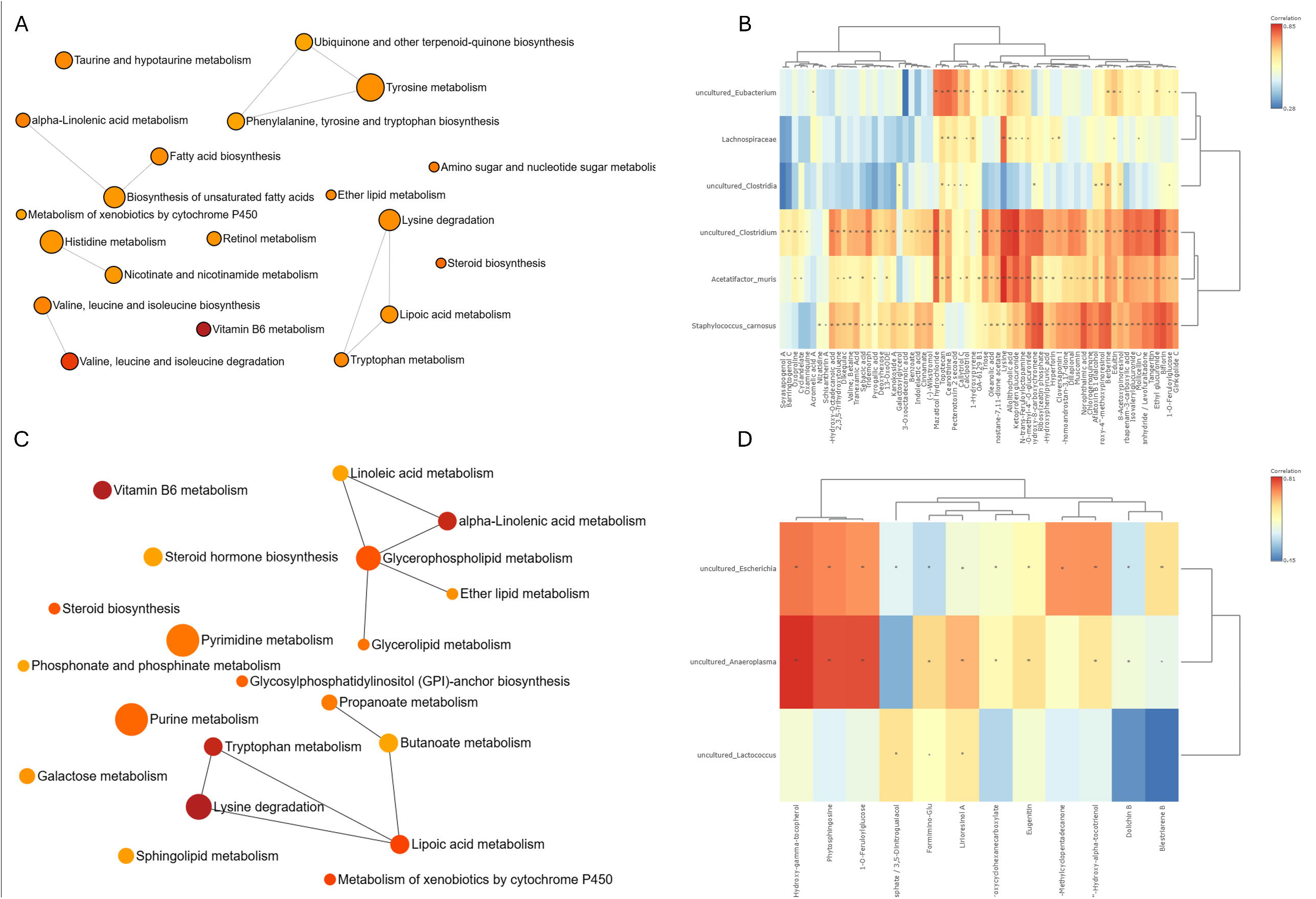
Integrated microbiome-metabolome analyses identify sex-associated microbial-metabolic networks during BA-modulated ARDS. (A) Integrated pathway network analysis of female BA+SEB stool datasets showing associations among pathways involved in amino acid metabolism, lipid remodeling, bile acid-associated signaling, oxidative stress, and related metabolic processes. Node size and other visual attributes are defined according to the pathway-analysis output shown in the panel. (B) Correlation heatmap showing associations between female stool bacterial taxa and metabolite features. Broad associations were observed for taxa including uncultured_Clostridium, Acetatifactor muris, and Staphylococcus carnosus. (C) Integrated pathway network analysis associated with male BA+SEB metabolomic profiles, including pathways related to nucleotide, lipid, amino acid, sphingolipid, and short-chain fatty acid metabolism. (D) Correlation heatmap of stool microbial taxa and BALF metabolite features in male BA+SEB mice. Associations involving uncultured_Escherichia, uncultured_Anaeroplasma, and uncultured_Lactococcus are shown. Heatmap colors indicate the strength and direction of Spearman correlation coefficients.

In males, integrated analyses identified a different metabolic network characterized by pathways related to nucleotide, lipid, amino acid, sphingolipid, and short-chain fatty acid metabolism (**Figure 7C**). Correlation analysis revealed distinct associations involving uncultured *Escherichia*, uncultured *Anaeroplasma*, and uncultured *Lactococcus* with BALF metabolite features (**Figure 7D**).

Together, these analyses demonstrate that BA-associated microbial and metabolic relationships differ between female and male mice during SEB-induced ARDS, supporting the presence of sex-specific gut-lung immunometabolic states.

## Discussion

### BA Colonization Drives Sex-Specific Gut-Lung Microbiome-Metabolome Remodeling During ARDS

The present findings demonstrate that *Bacteroides acidifaciens* (BA) colonization is associated with distinct sex-dependent microbial and metabolic responses during SEB-induced ARDS. In females, BA-associated disease exacerbation occurred alongside intestinal microbiome restructuring and metabolic remodeling involving inflammatory lipids, bile acid-associated pathways, and redox homeostasis. In contrast, the protective phenotype observed in males was accompanied by a distinct microbial community and metabolic profile characterized by short-chain fatty acid-associated signaling, lipid remodeling, and changes in amino acid, redox, and nucleotide metabolism. Direct comparisons further showed that female and male BA+SEB mice occupy distinguishable intestinal microbial and metabolic states.

These findings extend growing evidence that ARDS is influenced by gut-derived immune and metabolic signals rather than representing an exclusively pulmonary disorder [3-5]. Although both sexes received the same BA exposure, their microbial communities and stool and pulmonary metabolomic profiles differed, suggesting that host sex modifies the biological context in which BA interacts with the host. These observations build upon our previous study showing that BA exacerbated SEB-induced ARDS in females while attenuating disease in males [9]. In that study, the divergent responses were accompanied by differences in pulmonary inflammation, epithelial barrier integrity, immune-cell responses, oxidative stress, and mucosal immunity, and inhibition of estrogen synthesis partially attenuated the female phenotype [9].

Together, the current and previous findings support a working model in which host sex, and potentially the hormonal environment, influences the microbial and metabolic consequences of BA colonization during inflammatory lung injury. However, the relationships identified here are primarily associative; therefore, the microbial taxa and metabolic pathways identified in this study should be considered candidate mechanisms rather than established drivers of the divergent ARDS phenotypes.

### Distinct Immunometabolic Programs Accompany BA-Mediated Disease Exacerbation in Females and Protection in Males

Direct comparison of female and male BA+SEB mice demonstrated that the opposing effects of BA occur within qualitatively distinct metabolic environments rather than reflecting different magnitudes of a shared response. In females, BA-associated disease exacerbation was accompanied by coordinated remodeling of inflammatory lipid, bile acid, redox, and amino acid-associated pathways. In contrast, males exhibited a metabolic profile characterized by increased short-chain fatty acid-associated signaling, sphingolipid remodeling, and alterations in amino acid, redox, and nucleotide metabolism. These differences parallel the divergent inflammatory phenotypes established in our previous study and identify candidate metabolic processes that may contribute to sex-dependent responses to BA [9].

One of the most prominent features of the female metabolic profile was enrichment of leukotriene- and oxylipid-associated metabolites, including leukotriene B4 (LTB4), across the direct sex comparisons. Lipid mediators can regulate leukocyte recruitment and inflammatory signaling, and pulmonary eicosanoid responses can differ according to sex [16]. Oxidized lipid products can also influence macrophage activation and reactive oxygen species production [17,18]. Their enrichment in female BA+SEB mice therefore provides a potential metabolic link to the enhanced neutrophilic inflammation and oxidative stress previously observed in these animals [9]. Alterations in bile acid and redox-associated pathways may further contribute to this inflammatory environment. Bile acids can influence epithelial and immune function through receptors including FXR and TGR5 [10,19,20], while disruption of redox homeostasis can amplify inflammatory signaling and tissue injury [15,17,18]. Together, these findings suggest that BA-associated disease exacerbation in females occurs within a metabolic environment enriched in pathways capable of reinforcing inflammation, although the present data do not establish that these metabolic alterations directly drive disease severity.

The male response was distinct rather than uniformly opposite. Increased stool butyrate was particularly notable because short-chain fatty acids support epithelial barrier integrity, immune regulation, and communication along the gut-lung axis [12,13]. Butyrate can also promote colonic regulatory T-cell differentiation [21]. This finding is consistent with our previous observation that BA improved intestinal barrier integrity and reduced pulmonary inflammation in male mice [9]. Male BA+SEB mice also exhibited remodeling of sphingolipid and unsaturated fatty acid pathways, including reduced dihydroceramide. Because sphingolipid metabolites regulate membrane homeostasis, cellular stress, apoptosis, and inflammatory signaling [11,22], and sex-dependent differences in pulmonary sphingolipid biology have been reported [23], these changes are more appropriately interpreted as adaptive lipid remodeling than as a globally anti-inflammatory metabolic shift. Alterations in pulmonary redox, amino acid, and nucleotide metabolism further support the presence of a metabolically distinct male response associated with reduced inflammatory injury.

Importantly, sex-associated metabolic differences were not uniform across biological compartments. Some features, including LTB4, showed concordant female-biased abundance in stool and BALF, whereas others differed in magnitude or direction between compartments. This argues against a model in which intestinal metabolic changes are simply reproduced in the lung. Instead, BA-associated remodeling may influence pulmonary immunity through multiple mechanisms, including microbial metabolite production, modification of host-derived metabolites, intestinal barrier regulation, systemic immune signaling, and secondary tissue-specific metabolic responses [3-5]. Accordingly, the present findings support coordinated gut-lung metabolic remodeling but do not demonstrate direct transport of individual metabolites from the intestine to the pulmonary compartment.

### Integrated Microbiome-Metabolome Networks Reveal Sex-Associated Host-Microbial States

Integrated microbiome-metabolome analysis further demonstrated that BA-associated metabolic remodeling occurred within distinct microbial ecological contexts in females and males. In females, broader taxon-metabolite associations involved *Clostridium*-associated taxa, *Acetatifactor muris*, and *Staphylococcus carnosus* and converged with lipid-, bile acid-, amino acid-, and redox-associated metabolic pathways. In males, a compositionally distinct and more restricted network involved uncultured *Escherichia, Anaeroplasma*, and *Lactococcus*, with associations spanning sphingolipid, nucleotide, amino acid, and short-chain fatty acid-associated metabolism. These contrasting network structures reinforce the conclusion that the divergent responses to BA are accompanied by distinct microbial-metabolic configurations rather than a uniform microbiome response that differs only in magnitude.

The female network is particularly consistent with the lipid-centered phenotype identified independently by metabolomic profiling. Associations involving lipid- and fatty acid-derived metabolites occurred alongside the enrichment of leukotriene- and oxylipid-associated features observed in female BA+SEB mice. In males, associations involving metabolites such as phytosphingosine similarly converged with the independent identification of sphingolipid pathway remodeling. Integrating microbial-community and metabolomic data can identify coordinated ecological and metabolic patterns that are not apparent from either dataset alone [24-26].

These correlations, however, should not be interpreted as evidence that the associated bacterial taxa directly produce or regulate the corresponding metabolites. Statistical associations between microbiome and metabolome features may reflect direct microbial metabolism, host modification of microbial products, or shared biological responses rather than direct biosynthetic relationships [24-26]. Metabolic cross-feeding among community members provides an additional mechanism through which the abundance of one organism may be associated with metabolites produced or consumed by other community members [27]. Thus, the integrated analysis is most useful for identifying coordinated microbial-metabolic relationships and prioritizing candidate pathways for functional testing rather than establishing causal interactions.

### An Integrated Model for Sex-Dependent BA-Mediated Modulation of ARDS

Taken together, our previous and current findings support a working model in which host sex shapes the biological consequences of *Bacteroides acidifaciens* colonization through coordinated interactions among the hormonal environment, intestinal microbial community, host metabolism, and pulmonary immune response. Our previous study established that BA exacerbates SEB-induced ARDS in females while attenuating disease in males and further showed that inhibition of estrogen synthesis with letrozole partially improves the female BA+SEB phenotype [9]. The present study extends those findings by demonstrating that these divergent inflammatory outcomes are accompanied by distinct intestinal microbial communities, stool and pulmonary metabolomic profiles, and taxon-metabolite associations.

In females, BA-associated disease exacerbation occurs within an immunometabolic environment characterized by inflammatory lipid and oxylipid remodeling, altered bile acid and redox-associated metabolism, and broader microbial-metabolic network connectivity. These features align with the increased neutrophilic and Th17-associated inflammation, oxidative stress, barrier dysfunction, and IgA responses observed previously. In males, BA colonization is associated with a distinct metabolic state involving increased butyrate, sphingolipid remodeling, and alterations in amino acid, redox, and nucleotide metabolism, corresponding with improved epithelial barrier integrity, increased IL-22, and reduced pulmonary inflammation [9]. Thus, the male response appears to represent an alternative immunometabolic program rather than a weaker version of the female response.

The compartment-specific differences observed between stool and BALF further suggest that gut-lung communication is unlikely to reflect simple transfer of intestinal metabolites to the lung. Instead, BA-associated microbial remodeling may influence pulmonary immunity through a combination of microbial metabolite production, modification of host-derived metabolites, intestinal barrier effects, systemic immune signaling, and tissue-specific metabolic responses [3-5]. Within this framework, sex-dependent host factors, including the hormonal environment, may act upstream to shape how BA alters microbial ecology and metabolism. Bidirectional interactions between estrogen signaling and the gut microbiome have been described and provide biological precedent for such a model [28,29]. These relationships remain mechanistic hypotheses rather than established causal pathways, but they provide a framework for testing how sex modifies microbiome-dependent regulation of inflammatory lung injury.

### Limitations and Future Directions

Several limitations should be considered when interpreting these findings. First, the microbiome-metabolome relationships identified here are associative and do not establish causality. Taxon-metabolite correlations may reflect direct microbial metabolism, metabolic cross-feeding, host metabolism of microbial products, or parallel responses to inflammation [24-27]. Future studies using targeted microbial manipulation, gnotobiotic approaches, or defined microbial communities will be necessary to determine whether specific taxa contribute directly to the metabolic programs associated with BA-mediated protection or disease exacerbation.

Second, 16S rRNA sequencing provides limited species- and strain-level resolution, and several taxa contributing to the sex-associated networks could be classified only at higher taxonomic levels. Shotgun metagenomic or metatranscriptomic approaches could provide greater taxonomic and functional resolution and help determine whether differences in microbial community composition are accompanied by changes in metabolic capacity [24-26].

Interpretation of the metabolomic data is also limited by the putative annotation of many features. Untargeted metabolomics provides broad coverage of metabolic remodeling but does not definitively establish metabolite identity or absolute abundance. Targeted LC-MS/MS using authentic standards will therefore be important for validating candidate metabolites central to the proposed model, particularly inflammatory lipid mediators, bile acids, sphingolipid intermediates, tryptophan metabolites, and short-chain fatty acids [30].

The compartment-specific differences between stool and BALF also preclude conclusions regarding direct metabolite trafficking along the gut-lung axis. Future studies incorporating plasma metabolomics, longitudinal sampling, and stable-isotope tracing could distinguish direct movement of microbial metabolites from secondary host metabolic responses and clarify how intestinal, systemic, and pulmonary metabolic changes are temporally connected.

Finally, the mechanisms through which sex produces the divergent microbial and metabolic responses remain unresolved. Our previous finding that letrozole attenuated several components of the female BA-associated phenotype implicates the hormonal environment [9], but the relationship between estrogen signaling and the microbial-metabolic signatures identified here has not been established. Combining hormonal manipulation with longitudinal microbiome and metabolomic profiling will help determine whether sex hormones act upstream of microbial remodeling, through host metabolic and immune pathways, or at multiple levels [28,29].

Together, these future studies will enable functional testing of the candidate pathways identified here, including female-associated inflammatory lipid signaling and male-associated butyrate and sphingolipid metabolism. Such experiments will be necessary to distinguish biomarkers of the divergent BA responses from mechanisms that directly contribute to ARDS severity.

## List of Abbreviations

AhR: aryl hydrocarbon receptor
ARDS: acute respiratory distress syndrome
ASV: amplicon sequence variant
BA: *Bacteroides acidifaciens*
BALF: bronchoalveolar lavage fluid
CFU: colony-forming units
dCDP: deoxycytidine diphosphate
FDR: false discovery rate
FXR: farnesoid X receptor
IgA: immunoglobulin A
IL-17: interleukin-17
IL-22: interleukin-22
KEGG: Kyoto Encyclopedia of Genes and Genomes
LC-MS: liquid chromatography-mass spectrometry
LTB4: leukotriene B4
PCA: principal component analysis
PCoA: principal coordinates analysis
PBS: phosphate-buffered saline
PLS-DA: partial least squares discriminant analysis
ROS: reactive oxygen species
rRNA: ribosomal RNA
SCFA: short-chain fatty acid
SEB: staphylococcal enterotoxin B
SEM: standard error of the mean
TGR5: Takeda G protein-coupled receptor 5
Th17: T helper 17

## Ethics Approval

All animal experiments were conducted in accordance with the guidelines set forth by the National Institutes of Health Guide for the Care and Use of Laboratory Animals and were approved by the Institutional Animal Care and Use Committee (IACUC) of the University of South Carolina under protocol number 2687-101832-101923, approved October 19, 2023. No human subjects, human-derived materials, or patient data were used in this study.

## Author Contributions

SS: Conceptualization, Methodology, Investigation, Formal analysis, Visualization, and Writing-original draft. CC: Formal analysis, Visualization, and Review and Editing. SO: Investigation. JW: Investigation. AA: Writing-review and editing. KW: Conceptualization, Methodology, Formal analysis, Writing-review and editing, Resources, Supervision, and Funding acquisition.

## Acknowledgments

The authors would like to thank Dr. Brandon Busbee for donating the *Bacteroides acidifaciens* strains utilized in this study. The author gratefully acknowledges the CAM COBRE lead by Drs. Mitzi and Prakash Nagarkatti for providing a supportive academic environment and access to equipment utilized in this study. The author also acknowledges General Metabolics for performing the untargeted metabolomic analyses and providing technical support for metabolomic data acquisition and processing.

## Data Availability Statement

The datasets for this study can be found in the Figshare data repository at https://doi.org/10.6084/m9.figshare.34028607.

## Figure Legends

**Supplementary Figure S1. BA colonization is associated with differential metabolomic profiles in female stool and BALF during SEB-induced ARDS**.

(A) Hierarchical clustering heatmap of stool metabolite features in female SEB and BA+SEB mice. (B) Volcano plot showing stool metabolite differences between female BA+SEB and SEB groups. (C) Hierarchical clustering heatmap of BALF metabolite features in female SEB and BA+SEB mice. (D) Volcano plot showing BALF metabolite differences between female BA+SEB and SEB groups. Heatmaps display normalized metabolite abundance as indicated by the corresponding scale; selected metabolite features are labeled in volcano plots.

**Supplementary Figure S2. BA colonization is associated with differential metabolomic profiles in male stool and BALF during SEB-induced ARDS**.

(A) Hierarchical clustering heatmap of stool metabolite features in male SEB and BA+SEB mice. (B) Volcano plot showing stool metabolite differences between male BA+SEB and SEB groups. (C) Hierarchical clustering heatmap of BALF metabolite features in male SEB and BA+SEB mice. (D) Volcano plot showing BALF metabolite differences between male BA+SEB and SEB groups. Heatmaps display normalized metabolite abundance as indicated by the corresponding scale; selected metabolite features are labeled in volcano plots.

**Supplementary Figure S3. Relative abundance of selected bacterial taxa across treatment groups in female and male mice**.

Relative abundance of selected stool bacterial taxa is shown across VEH, BA, SEB, and BA+SEB groups in (A) female and (B) male mice. Individual points represent biological samples, with summary statistics displayed as indicated in the figure. Selected taxa were chosen to further illustrate treatment-associated abundance patterns observed in the microbiome dataset.

**Supplementary Figure S4. Taxonomic abundance and prevalence patterns across experimental groups in female and male stool microbiomes**.

(A) Taxonomic abundance patterns across VEH, BA, SEB, and BA+SEB groups in female mice. (B) Prevalence of bacterial taxa across female treatment groups. (C) Taxonomic abundance patterns across treatment groups in male mice. (D) Prevalence of bacterial taxa across male treatment groups. Prevalence represents the proportion of samples in which individual taxa were detected.

**Supplementary Figure S5. Relative abundance of intestinal bacterial genera in BA and BA+SEB mice**. Relative abundance of dominant bacterial genera in (A) female and (B) male stool microbiomes following BA colonization alone or BA colonization followed by SEB challenge. Stacked bars represent the proportional contribution of the displayed genera to each microbial community, with remaining taxa grouped as “Other.”

