## Supplementary figures and images for "Sex-Dependent Microbiome-Metabolome Remodeling Associated with *Bacteroides acidifaciens* Modulation of Acute Respiratory Distress Syndrome"

### Supplemental Figure 1

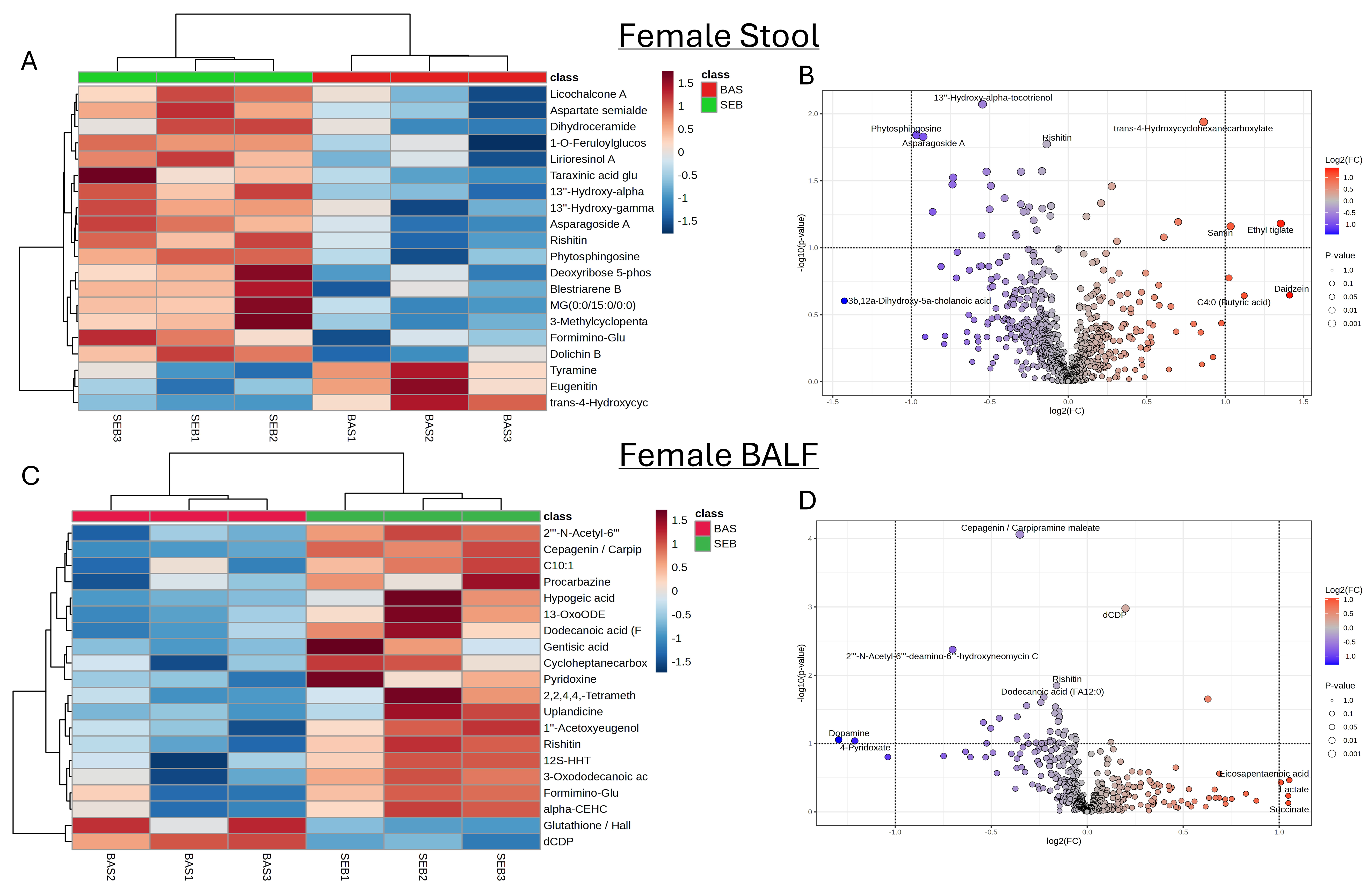

### Supplemental Figure 2

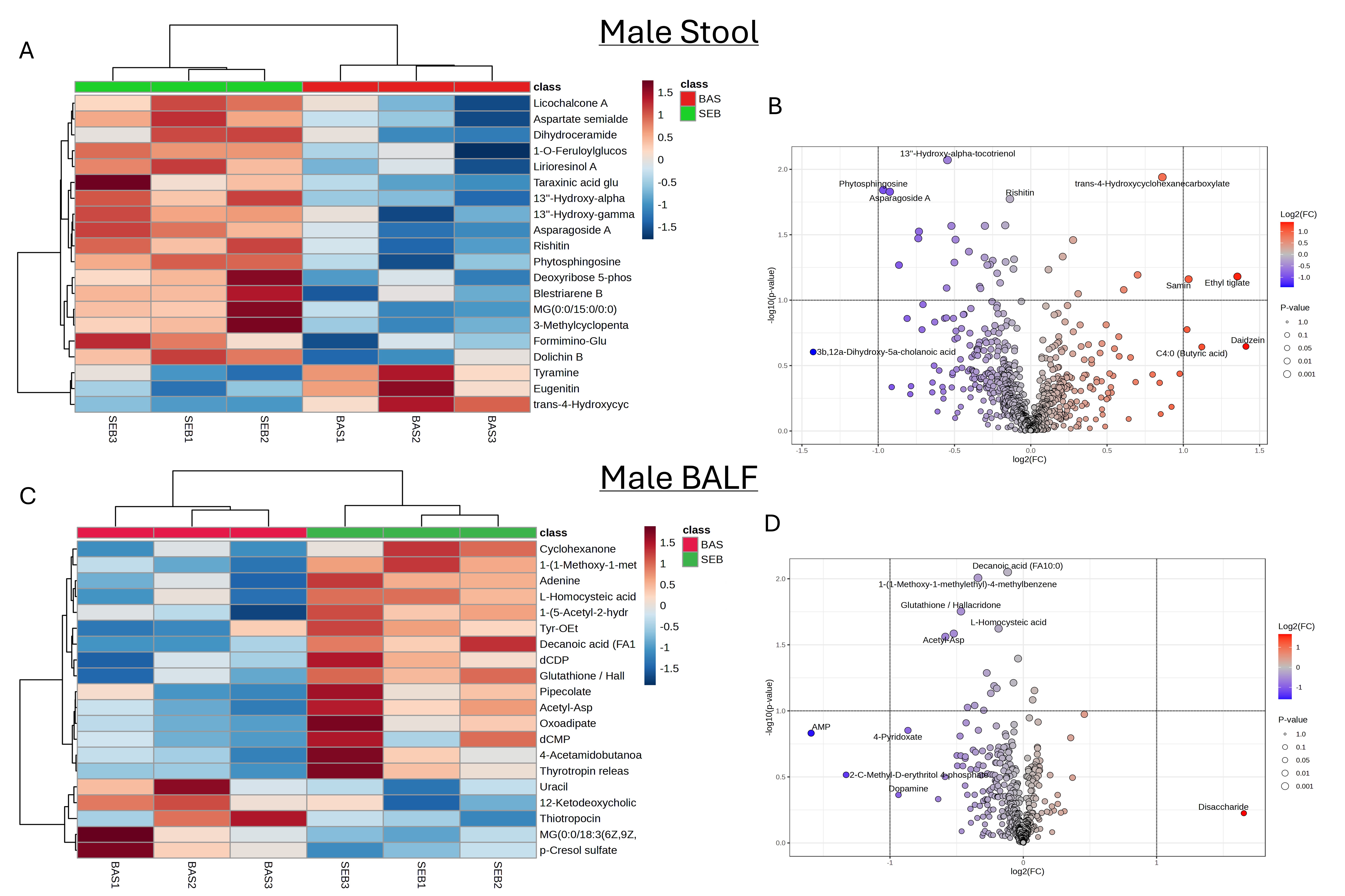

### Supplemental Figure 3

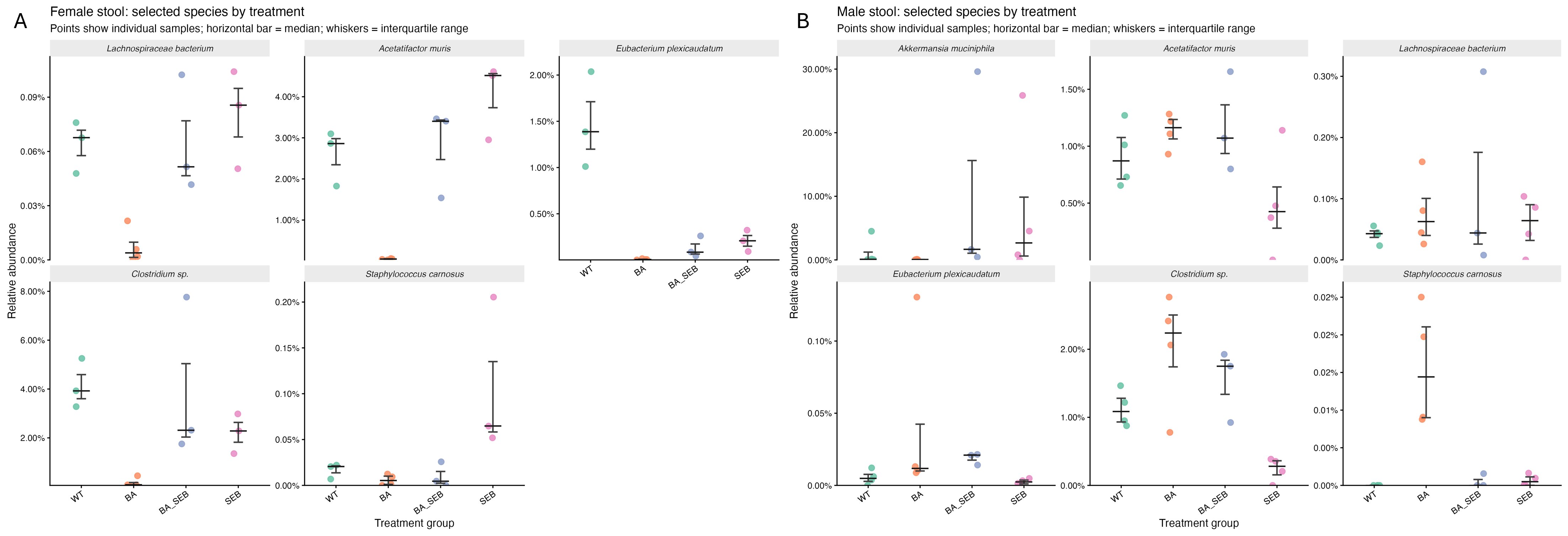
